# MiGenPro: A linked data workflow for phenotype-genotype prediction of microbial traits using machine learning

**DOI:** 10.1101/2025.08.21.671437

**Authors:** Mike Loomans, Maria Suarez-Diez, Peter J. Schaap, Edoardo Saccenti, Jasper J. Koehorst

**Affiliations:** Laboratory of Systems and Synthetic Biology, Wageningen University & Research, Wageningen, The Netherlands; UNLOCK, Wageningen University & Research & Research and Delft University of Technology, The Netherlands

**Keywords:** Bioinformatics, Machine Learning, Genomics (microbiology), Visualization, Python library, Linked data, Genotype-phenotype association

## Abstract

The availability of microbial genomic data and the development of machine learning methods have created a unique opportunity to establish associations between genetic information and phenotypes. Here, we introduce a computational workflow for Microbial Genome Prospecting (MiGenPro) that combines phenotypic and genomic information. MiGenPro serves as a workflow for the training of machine learning models that predict microbial traits from genomes that have been annotated. Microbial genomes have been consistently annotated and features were stored in a semantic framework that is easy to query using SPARQL. The data was used to train machine learning models and successfully predicted microbial traits such as motility, Gram stain, oxygen requirement, optimal temperature range, and sporulation capabilities. To ensure robustness, a hyperparameter halving grid search was used to determine optimal parameter settings followed by five randomised train-test split validations which demonstrated consistent model performance across iterations and without overfitting. Effectiveness was further validated through comparison with existing models, showing comparable accuracy, with modest variations attributed to differences in datasets rather than methodology. Classification can be further explored using feature importance characterisation to identify biologically relevant genomic features. MiGenPro provides an easy to use interoperable workflow to build and validate models to predict phenotypes from microbes based on their annotated genome.

**Key Messages:**

- MiGenPro merges phenotypic and genomic data for microbial trait prediction using machine learning.
- MiGenPro mitigates phenotype prediction model constraints by leveraging linked data technologies.
- MiGenPro’s FAIR design enables adaptation for various phenotypes with available training data.

## Introduction

Advances in experimental and computational techniques for DNA extraction, sequencing, assembly, and structural and functional annotation have increased the availability of microbial genotypes. As a result of the increasing throughput of these approaches, millions of genome sequences are stored in databases using highly structured formats that collect both genetic sequences and the result of annotations that inform on genomic features. The potential to generate and store sequence data dwarfs the availability of phenotype information. The characterization of the microbial phenotypes linked to sequences still lags behind as dedicated experiments are often required to characterise each aspect such as optimal temperature range, substrate utilisation potential or potential for sporulation among others.

Yet the exploitation of microbes depends on the microbial phenotype. Consider the global biotechnology sector that is expanding in the coming years(23). The expansion of this industry is coupled with developments in strain optimization and the discovery of strains with industrial potential(6; 2). The selection of industrial strains often considers an increased performance of the cell factory and relevant traits include, alongside desired metabolic pathways, thermotolerance, osmotolerance, and general robustness(27). For example, the production of polyhydroxyalkanoates in *Schlegelella thermodepolymerans* benefits from high temperatures as it reduces the risk of contamination(22). Therefore, exploring these phenotypes can accelerate biotechnological progress. Similarly, bioremediation strategies could benefit from accurate prediction of phenotypes to aid the development of microbial communities that promote soil health.

These applications demonstrate the potential of methods to accurately predict microbial phenotypes from genome sequences. The use of public genome repositories and annotation pipelines to predict prokaryotic phenotypes has been widely practiced(24; 33; 12; 19). However, these approaches are challenged by the lack of consistently annotated genomes in a format that supports automated querying and recovery of strain associated phenotypes. Additionally, these methods also struggle with species-specific genetic elements as these limit the resolution of the prediction tools.

Recently, the work of Koblitz, et al. 2025 illustrates the potential of machine learning approaches when applied to the task of predicting phenotype information from genome data(19). It also highlights the need for uniform data annotation and preserving the association between genotype and phenotype data in the training datasets. Here, we demonstrate how to combine semantic technologies for data storage and linking and machine learning models to predict phenotypes from genome data. Currently relevant and publicly available databases in the life sciences provide the means to retrieve their data using machine readable data formats (such as JSON). Here, we provide a workflow to exploit this technology that can be adapted and extended to any relevant phenotype for which training data are available. The approach can also be used to test the performance of alternative prediction approaches. The Microbial Genome Prospecting (MiGenPro) phenotype prediction workflow encapsulates the steps required to go from a collection of genome sequences and phenotype information to machine learning models.

These steps include data retrieval using REST API, the use of the Genome Biology Ontology Language (GBOL) together with SPARQL and RDF for data storage, and standardised workflows for uniform genome data annotation(10; 1). We have showcased the effectiveness of the MiGenPro workflow with the prediction of physiological characteristics using various machine learning models trained on data automatically retrieved from the BacDive repository(31). The predictions are complimented with the addition of methods for feature selection that extract biological knowledge from the developed models.

## Methods

The modules in the MiGenPro workflow contribute to two main tasks, data formatting and prediction of phenotypes, as illustrated in Figure 1. The modular design ensures that any step can be run independently provided that the input data is available.

**Figure 1.**
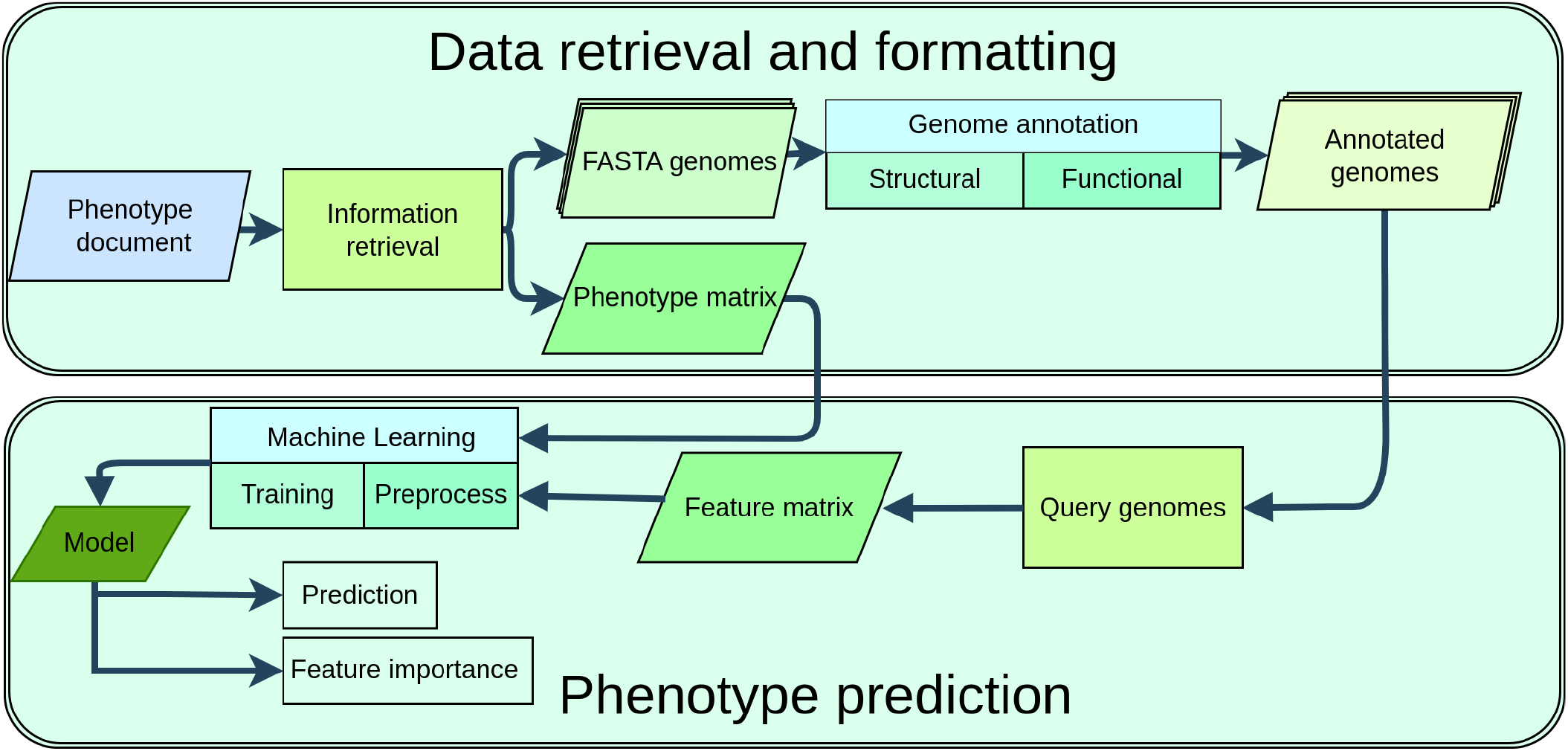
The workflow starts with by loading a phenotype graph, which is queried using SPARQL to generate a list of genome IDs and a matrix containing the phenotypes. These IDs are used to obtain corresponding FASTA files and the phenotype matrix links each genome ID to its respective phenotype. The FASTA genomes are annotated with SAPP (Semantic Annotation Platform with Provenance) that runs InterProScan and prodigal and outputs them in GBOL format. The annotated genomes are queried with SPARQL to extract genomic features. The phenotype and feature matrices are loaded into Python where a series of classes prepare the data and train the models. The resulting model can predict phenotypes, and biological relevance is determined by assessing feature importance.

### Retrieval of phenotype graph

The term phenotype graph refers to a document that describes the data available in the database(s) from which data are to be retrieved. This document should contain information on genome identifiers and respective phenotypes. Here we retrieved the phenotype graph containing Genome identifiers alongside selected phenotypes namely, Gram stain, motility, oxygen requirement, optimal growth temperature, and spore formation from the BacDive database. The BacDive database REST API was used to retrieve this information in JSON format. The JSON results were transformed into a linked data structure, JSON-LD, using SAPP (v2.0)(21).

### Information retrieval: Query phenotypes and genomes

The linked data structure was converted into a HDT (Header Dictionary Triple) file using SAPP (v2.0) and subsequently queried using the SPARQL integration within SAPP using the queries in supplementary file 1(1). The phenotype data were parsed into a columnar format using Python (v3.12.5) with the pandas library (v2.1.1), creating a matrix. This matrix contained for each phenotype a set of rows denoting the various genomes with NCBI genome identifiers and columns for the features. The matrix was filtered and only physiological characteristics for which at least 500 annotated genomes were available were considered.

For species for which multiple genomes were available, the ete4 (v.3.1.3) library was used for its NCBI taxonomy tree to ensure *≤*10 randomly chosen genomes per species to avoid over-representation of model organisms(16). The genome data in FASTA format was retrieved using the NCBI Datasets conda package(28).

### Genome annotation

The genome data were then structurally and functionally annotated using a standardised workflow written in the Common Workflow Language(9; 20). This workflow contains the following SAPP modules: Genome conversion to RDF, Prodigal (v2.6.3) for gene prediction and InterProScan (v5.53-87.0) for functional annotation (21; 18; 3). The annotated results are represented in RDF following the GBOL (Genome Biology Ontology Language) structure using the HDT format. Each genome is encapsulated in a single HDT file.

## Phenotype prediction

### Query genomes and feature matrix construction

The protein domain information was extracted from each genome separately using SPARQL queries (supplementary file 2). The resulting protein domains were merged into a single matrix containing all protein domain frequencies across the genomes considered for the selected phenotype.

For each phenotype, feature selection was performed on the training data only, after the train-test split, to avoid information leakage into the held-out test set. Selection proceeded in two steps. First, a variance threshold filter removed protein domains with low variance across the training genomes to discard uninformative features. Second, the scikit-learn mutual info classif function was used to calculate the mutual information between each remaining protein domain and the phenotype, and the top 25% of protein domains ranked by mutual information were retained. This reduced the dimensionality of the dataset. The selected protein domains were then used as features for the machine learning models, and the same selection was applied to the test data.

### Machine learning data preprocessing

The phenotype and feature matrices were used as starting point for training machine learning models. The dataset was split into an 80% train and 20% test with the scikit-learn train test split function(29). The train test split function randomly splits the data into two subsets, ensuring that the distribution of the target variable is preserved in both the training and test sets with the stratify flag. Training data was oversampled using SMOTEN (Synthetic Minority Over-sampling Technique for Nominal) oversampling from the imbalanced-learn library (v0.14.2) (7). This technique was used for categorical oversampling with a maximum oversampling factor of two to balance the difference in frequency amongst phenotypes.

## Machine Learning training

Machine learning methods based on decision trees were selected due to their interpretability. The algorithms, random forest, decision trees, and gradient boosting were imported from the scikit-learn (v1.5.2) library in Python (29). XGBoost was imported from the xgboost library (v2.1.1) (8).

Parameter optimisation was performed per phenotype and classifier using the HalvingGridSearchCV function. To control overfitting, the grid space was constrained to favour shallower trees and higher minimum sample thresholds. Decision tree models explored maximum tree depths of 3, 5, and 10 with minimum samples per leaf of 5, 10, and 20. Random forest models tuned the number of trees (100, 200, 500, 1000), maximum depth (5, 10, 15), minimum samples per leaf (5, 10, 20, 50), and minimum samples per split (5, 10, 20). Gradient boosting models explored learning rates (0.01, 0.05, 0.1), maximum depth (3, 5), minimum samples per split (5, 10, 20, 50), subsampling ratio (0.7, 0.8, 1.0), and estimator counts (50, 100, 200, 500). XGBoost models additionally explored L2 regularisation (1.0, 5.0, 10.0), column subsampling (0.7, 0.8), and minimum child weight (1, 5, 10).

The parameter max resources was set to the number of samples and min resources was set to exhaust with the Matthews correlation coefficient as the target metric. A successive halving grid search first trains all parameter combinations with few resources, in this scenario a small subset of data; it then selects the best performing parameter combinations and repeats the process with more resources until the optimal combination has been found.

The HalvingGridSearchCV selected the following regularised configurations for each phenotype. Decision tree classifiers used maximum depths of 5–10, minimum samples per leaf of 5– 10, and minimum samples per split of 10–50. Random forest classifiers consistently selected a maximum depth of 15 and a minimum of 5 samples per leaf (5); the number of trees ranged from 200 (temperature) to 1000 (Gram stain, motility, spore formation), with minimum samples per split of 5–20. Gradient boosting classifiers used learning rates of 0.05–0.1, maximum depths of 3–5, minimum samples per split of 5– 50, subsampling ratios of 0.7–0.8, and 500 estimators (13). XGBoost classifiers used a learning rate of 0.1, maximum depths of 5–7, subsampling ratios of 0.7–0.8, column subsampling of 0.7–0.8, a minimum child weight of 1, L2 regularisation (*λ* = 1.0–10.0), and 200–500 estimators (8).

### Model assessment and feature importance

The models were evaluated in accuracy, Matthews correlation coefficient, macro-averaged F1 score, and AUC on a held-out test set which consisted of 20% of the data. Performance metrics were visualised using matplotlib (v3.8.2). The models’ feature importance was inferred using average Gini values from five train test cycle validations with the scikit learn library. The feature importance results of each model were then ordered by their cumulative rank sum. For each iteration the Gini scores were ordered and then ranked, and these ranks were subsequently summed across iterations; the overall importance of each feature was then inferred from these summed ranks, with the lowest sums corresponding to the most important features.

## Results

The MiGenPro workflow offers a modular approach to building machine learning models that predict microbial phenotypes from genomic data. By integrating various linked data tools and techniques, the workflow streamlines the process from data retrieval and integration to model training and evaluation. A key advantage of leveraging linked data is the ability to seamlessly query public SPARQL endpoints to access experimentally validated phenotypic metadata from resources such as BacDive(31). This interoperability significantly reduces manual preprocessing and fosters the reuse of high-quality, curated data across studies.

### Performance of trained models

Model performance for decision tree, random forest, gradient boosting, and XGBoost models is shown in Table 2. Comparison to different studies shows that models reach similar bottlenecks in terms of performance due to the datasets being used. It should be noted that there are big differences in the training and testing datasets used for each study, and the results were directly taken from the respective studies. The differences between datasets used are vast, as Feldbauer, et al. 2015 used 97 genomes for training, Weimann, et al. 2016 used 234, Lingner et al. 2010 used 795 finished and 237 unfinished genomes, and Koblitz, et al. 2025 used between 3000 and 12000 genomes for training depending on the phenotype.

An overview of the model performance scores can be found in Table 3. Performance indicators for all methods and all physiological characteristics considered are high, indicating the suitability of this approach. The standard deviations observed from five random train-test splits remain low, indicating stable outcomes. Among the four model types tested, decision tree classifiers consistently underperform compared to the more sophisticated ensemble methods. XGBoost consistently achieves the highest scores, followed closely by gradient boosting and random forest. The decision tree trails the ensemble methods most markedly for oxygen requirement, where the gap exceeds 20% in macro-averaged F1 score. Motility and spore formation remain the most challenging phenotypes overall, as reflected in their comparatively low Matthews correlation coefficients.

### Feature analysis and biological interpretation

Tree-based methods can be analysed using Gini importance to evaluate feature importance. The Gini index evaluates how much each feature contributes to the performance of the classifier and can be used to select key features associated to the classification.

As an example, the 5 most relevant protein domains for predicting the motile potential are provided in Table 4.

The rank sum method aggregates the importance of the features across the five iterations. Within each iteration the features are ordered by their Gini importance and assigned a rank, with rank one denoting the most important feature, and these ranks are then summed across iterations. Lower total scores indicate higher overall importance, because features that are consistently ranked as critical across iterations accumulate smaller sums; a feature ranked first in every iteration attains the minimum possible sum of five.

In the analysis of protein domains to predict motility using a gradient boosting classifier, several domains consistently emerged with a low cumulative sum. Functional annotations and descriptions for the Pfam protein domain accessions were retrieved from InterPro and the primary literature cited in Table 4.

The PF02120 domain was identified as the most important domain, as shown by the rank sum of 5 over five iterations. This domain is located in the C-terminal of the FliK protein and controls flagellar hook length(32). The second most important domain, PF02203, is a ligand-binding domain of methyl-accepting chemotaxis receptors, while the third, PF00672 (HAMP), is likewise common in chemoreceptors(26; 17). These domains are associated with chemoreceptors and possibly chemotaxis, which determines the direction of motility. The fourth most important domain, PF06429, is the C-terminal domain shared by the flagellar basal-body rod and hook proteins (FlgE, FlgF, FlgG), core structural components of the flagellum(15). The fifth most important domain, PF00563, is associated with signal proteins containing the EAL domain that have been associated with regulation of motility(14; 11).

## Discussion

MiGenPro enables data curation and machine learning approaches for microbial traits. Previous work has illustrated the potential of machine learning models on high-quality curated datasets to predict bacterial phenotypes(24; 33; 12; 19). The application of this workflow requires genomes combined with a database or matrix with phenotype information of the corresponding genomes.

Consistent annotation of genome data prior to comparative genomic studies has been regarded as essential to prevent bias and artifacts due to differences in the tools used for genome annotation (30). MiGenPro enables automation of these tasks that facilitate comparative genomics. Here we have illustrated the use of MiGenPro using phenotype data from BacDive, and due to its modular design can be extended to other data resources using Linked Data formats. The starting point is a phenotype graph file that associates physiological measurements to standardised genome identifiers such as the ones in NCBI or an internal system. Once this starting point is available, the workflow provides the tools to retrieve and canonicalise the data and enable the development of machine learning models. By standardising genomic data as Linked Data, a high level of interoperability is obtained. This interoperability allows for efficient development of phenotype prediction learning models. The quality of the genomes was not compared to model performance; however, it stands to reason that genomes of higher-quality would yield more accurate results, as machine learning methods could use the absence of a protein domain to infer a phenotype(4).

The effectiveness of this workflow has been demonstrated by successful prediction of microbial traits: motility, Gram stain, oxygen requirement, optimal temperature range, and spore formation. Predictions derived from genomic features illustrate the workflow’s ability to handle complex biological data and provide insights.

### Model performance is not artificially inflated

Phenotypic traits often relate to the species and genus of microbes; however, exceptions are known where traits occur only in specific strains within a species or genus(30). When training classifiers for phenotype prediction, it is important that performance reflects genuine genotype-phenotype associations rather than memorisation of taxa-specific features. To this end, regularised hyperparameters were used: all classifiers used constrained tree depths, elevated minimum sample thresholds, and the boosting models applied subsampling and L2 regularisation. These constraints reduce the gap between training and test accuracy to at most a few percentage points, indicating that the models generalise rather than overfit the training data. The feature importances are highly stable across the five independent train-test iterations (Table 4), and the performance metrics show low standard deviations across these iterations (Table 3), indicating that the models do not depend on artefacts of a particular random split.

The biological coherence of the highest-ranking features further supports the validity of the predictions. As detailed in Table 4, the most important protein domains for motility map to well-characterised flagellar and chemotaxis machinery, consistent with the underlying biology. This interpretability indicates that the models learn relevant biological signal rather than fitting to noise, supporting the generalisability of the predictions to new, unseen genomes.

The Gini feature importance was used to assess the rank of these features. Gini is only applicable to tree-based methods. It should also be noted that Gini importance can be biased towards features with higher cardinality or variance, and the resulting rankings should be corroborated with complementary measures. The workflow is modular as the user could use other methods for the evaluation of feature importance, such as Shapley additive explanation values(25).

### Model performance matches previous phenotype prediction methods

Cross-study comparison of the best performing machine learning approaches in Table 2 reveals marginal performance variations (Δ accuracy *≤* 8%) barring motility prediction, suggesting that algorithm choice is less critical than data quality for phenotype prediction. From Table 3, the three ensemble methods (XGBoost, gradient boosting, and random forest) differ by at most 11% in macro-averaged F1 score per phenotype, with the largest spread occurring for temperature and oxygen requirement where class imbalance is most pronounced. The simpler decision tree classifier trails them, most markedly for oxygen requirement where the gap exceeds 20%.

The primary distinction between these studies lies in the datasets employed, thereby underscoring the need for a workflow capable of easily training models on new data. Comparing the methods themselves is challenging, as the accuracy metric is limited when evaluating model performance on imbalanced test datasets. This is particularly pertinent for the temperature and spore formation phenotypes, where the majority class (mesophilic; non-sporulating) dominates the dataset (Table 1); the high accuracy values (0.90–0.98) are therefore partly driven by majority-class agreement, and the Matthews correlation coefficient (Table 3) provides a more stringent and informative read of classifier performance on these imbalanced tasks. The accuracy metrics reported were directly sourced from the respective studies. For a comprehensive comparison of models developed throughout the last 15 years, accuracy was selected as one of the few performance metrics available(24; 33; 12; 19). A curiosity here is the case of the motility dataset that classifies all types of motility, gliding, swimming, twitching under the same header as being motile(31; 19). These motility subgroups are not evenly distributed and have varying levels of complexity, the heterogeneity in this phenotype results in lower than expected performance metrics.

**Table 1.** Number of genomes used per phenotype for training and testing models after filtering for a minimum of 500 genomes per phenotype and application of the 10 genomes per species cut-off.

| Phenotype | Count |
| --- | --- |
| **Gram** | 6391 |
| negative | 4127 |
| positive | 2264 |
| **Motility** | 5810 |
| no | 3254 |
| yes | 2556 |
| **Oxygen** | 9679 |
| aerobe | 4874 |
| anaerobe | 2715 |
| facultative anaerobe | 1064 |
| microaerophile | 1026 |
| **Spore** | 4263 |
| no | 3586 |
| yes | 677 |
| **Temperature** | 16204 |
| mesophilic | 15360 |
| thermophilic | 844 |

**Table 2.**
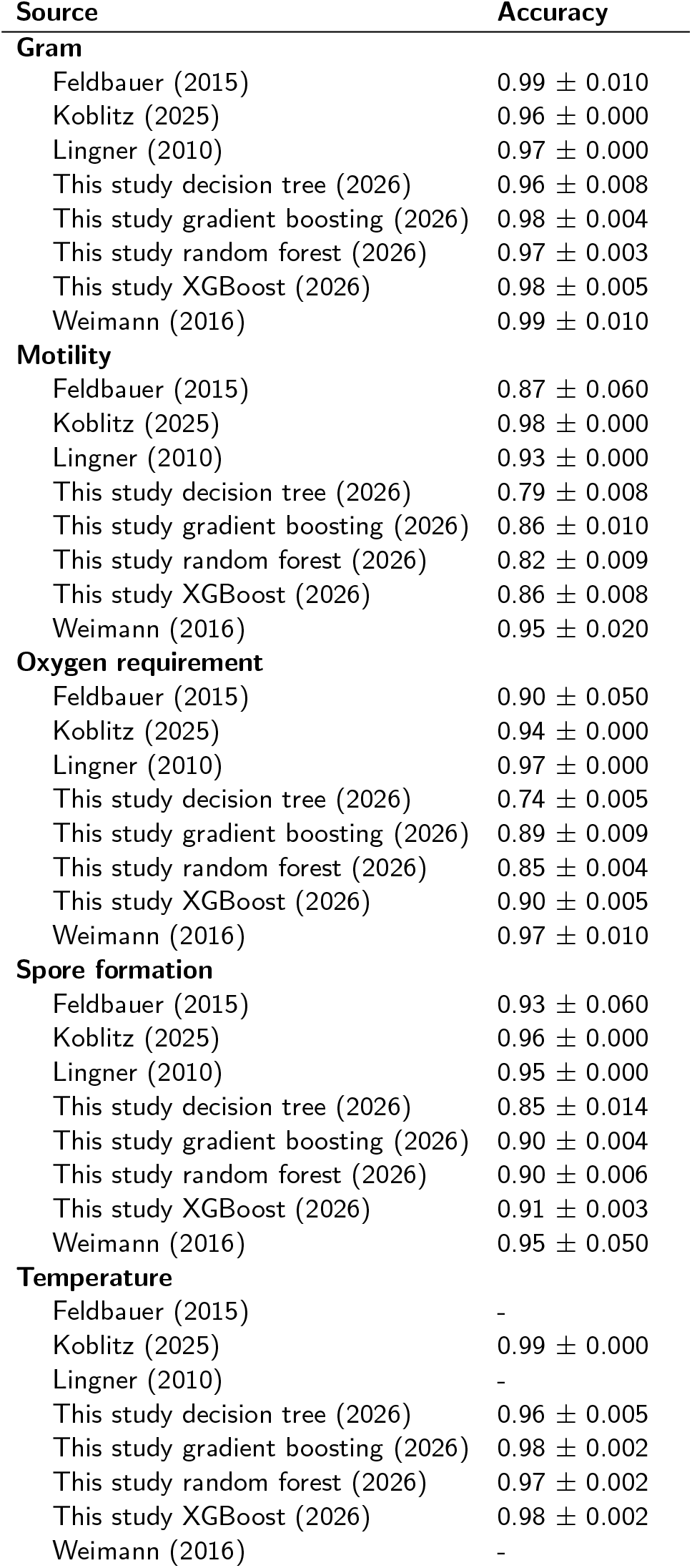
In-study reported accuracy metrics for the Gram stain, oxygen requirement, motility and spore formation phenotypes. Accuracy measurements were taken from the original studies and were not assessed using the same dataset. Temperature prediction was absent for studies other than Koblitz, et al. 2025 and values were rounded to two decimals. The mean and standard deviation are displayed in the table.

| Source | Accuracy |
| --- | --- |
| **Gram** |  |
| Feldbauer (2015) | 0.99 ± 0.010 |
| Koblitiz (2025) | 0.96 ± 0.000 |
| Lingner (2010) | 0.97 ± 0.000 |
| This study decision tree (2026) | 0.96 ± 0.008 |
| This study gradient boosting (2026) | 0.98 ± 0.004 |
| This study random forest (2026) | 0.97 ± 0.003 |
| This study XGBoost (2026) | 0.98 ± 0.005 |
| Weimann (2016) | 0.99 ± 0.010 |
| **Motility** |  |
| Feldbauer (2015) | 0.87 ± 0.060 |
| Koblitiz (2025) | 0.98 ± 0.000 |
| Lingner (2010) | 0.93 ± 0.000 |
| This study decision tree (2026) | 0.79 ± 0.008 |
| This study gradient boosting (2026) | 0.86 ± 0.010 |
| This study random forest (2026) | 0.82 ± 0.009 |
| This study XGBoost (2026) | 0.86 ± 0.008 |
| Weimann (2016) | 0.95 ± 0.020 |
| **Oxygen requirement** |  |
| Feldbauer (2015) | 0.90 ± 0.050 |
| Koblitiz (2025) | 0.94 ± 0.000 |
| Lingner (2010) | 0.97 ± 0.000 |
| This study decision tree (2026) | 0.74 ± 0.005 |
| This study gradient boosting (2026) | 0.89 ± 0.009 |
| This study random forest (2026) | 0.85 ± 0.004 |
| This study XGBoost (2026) | 0.90 ± 0.005 |
| Weimann (2016) | 0.97 ± 0.010 |
| **Spore formation** |  |
| Feldbauer (2015) | 0.93 ± 0.060 |
| Koblitiz (2025) | 0.96 ± 0.000 |
| Lingner (2010) | 0.95 ± 0.000 |
| This study decision tree (2026) | 0.85 ± 0.014 |
| This study gradient boosting (2026) | 0.90 ± 0.004 |
| This study random forest (2026) | 0.90 ± 0.006 |
| This study XGBoost (2026) | 0.91 ± 0.003 |
| Weimann (2016) | 0.95 ± 0.050 |
| **Temperature** |  |
| Feldbauer (2015) | - |
| Koblitiz (2025) | 0.99 ± 0.000 |
| Lingner (2010) | - |
| This study decision tree (2026) | 0.96 ± 0.005 |
| This study gradient boosting (2026) | 0.98 ± 0.002 |
| This study random forest (2026) | 0.97 ± 0.002 |
| This study XGBoost (2026) | 0.98 ± 0.002 |
| Weimann (2016) | - |

**Table 3.** Classifier performance for each phenotype using the macro-averaged F1 score, area under the receiver operating characteristic curve (AUC), and Matthews correlation coefficient (MCC) metrics to assess model performance. Mean and standard deviation values are derived from five train–test cycle validations of all models: DT (Decision Tree), GB (Gradient Boosting), RF (Random Forest), XGB (XGBoost).

| Classifier | F1<sub>macro</sub> | Matthews | AUC |
| --- | --- | --- | --- |
| **Gram** |  |  |  |
| DT | 0.958±0.008 | 0.916±0.017 | 0.973±0.006 |
| GB | 0.976±0.004 | 0.952±0.008 | 0.993±0.002 |
| RF | 0.970±0.003 | 0.942±0.006 | 0.994±0.001 |
| XGB | 0.979±0.005 | 0.957±0.010 | 0.996±0.002 |
| **Motility** |  |  |  |
| DT | 0.787±0.009 | 0.574±0.018 | 0.820±0.010 |
| GB | 0.854±0.010 | 0.708±0.021 | 0.924±0.007 |
| RF | 0.818±0.009 | 0.637±0.018 | 0.906±0.006 |
| XGB | 0.854±0.009 | 0.708±0.017 | 0.929±0.005 |
| **Oxygen requirement** |  |  |  |
| DT | 0.639±0.006 | 0.597±0.005 | 0.848±0.012 |
| GB | 0.832±0.013 | 0.828±0.014 | 0.964±0.004 |
| RF | 0.759±0.007 | 0.758±0.006 | 0.947±0.005 |
| XGB | 0.842±0.008 | 0.840±0.007 | 0.969±0.003 |
| **Spore formation** |  |  |  |
| DT | 0.711±0.016 | 0.424±0.033 | 0.760±0.031 |
| GB | 0.790±0.010 | 0.585±0.017 | 0.886±0.013 |
| RF | 0.765±0.014 | 0.557±0.028 | 0.901±0.017 |
| XGB | 0.812±0.011 | 0.632±0.017 | 0.913±0.014 |
| **Temperature** |  |  |  |
| DT | 0.772±0.023 | 0.555±0.048 | 0.788±0.019 |
| GB | 0.880±0.015 | 0.767±0.028 | 0.945±0.010 |
| RF | 0.779±0.018 | 0.622±0.028 | 0.945±0.009 |
| XGB | 0.890±0.012 | 0.788±0.025 | 0.955±0.009 |

**Table 4.** Gini feature importance for motility prediction with gradient boosting classifier visualised as the rank sum over five iterations.

| Domain | Rank sum | Description | Reference |
| --- | --- | --- | --- |
| PF02120 | 5 | C-terminal domain of FliK; controls flagellar hook length | Shibata, et al. 2007 (32) |
| PF02203 | 10 | Ligand-binding domain in methyl-accepting chemotaxis receptors | Mise 2016 (26) |
| PF00672 | 21 | HAMP domain common in chemoreceptors | Hulko, et al. 2006 (17) |
| PF06429 | 23 | C-terminal domain of the flagellar basal-body rod/hook proteins (FlgEFG) | Homma, et al. 1990 (15) |
| PF00563 | 23 | EAL domain found in signalling proteins | Galperin, et al. 2001 (14) |

### Impact of data quality on performance

BacDive data lacks or under-represent certain phenotypes (e.g., temperature ranges of extremophiles) as indicated by their omission in Table 1. Low-performance phenotypes such as motility that is derived from complex multi-gene regulation and have multiple methods for achieving the same result. The rapid development of AI technology represents a promising future area of research where phenotype analysis is based on gene sequences combined with protein structures to predict these more complex phenotypes. In addition, AI technology can also more easily process literature for the identification of historical records of phenotypes that are currently not incorporated into dedicated repositories such as BacDive(31).

## Abbreviations

AUC: Area Under the Curve
CWL: Common Workflow Language
DT: Decision Tree
FAIR: Findable, Accessible, Interoperable, Reusable
GB: Gradient Boosting
GBOL: Genome Biology Ontology Language
HDT: Header Dictionary Triples
JSON: JavaScript Object Notation
JSON-LD: JSON for Linking Data
MCC: Matthews correlation coefficient
ML: Machine Learning
MiGenPro: Microbial Genome Prospecting
NCBI: National Center for Biotechnology Information
REST API: Representational State Transfer Application Programming Interface
RDF: Resource Description Framework
RF: Random Forest
SAPP: Semantic Annotation Platform with Provenance
SMOTEN: Synthetic Minority Over-sampling Technique for Nominal
SPARQL: SPARQL Protocol RDF Query Language

## Data availability and description

The source code of MiGenPro is available free of use under the MIT license at https://gitlab.com/wurssb/migenpro. In addition, all data generated in this project, the annotated genomes are available as compressed and pre-indexed HDT files combined with the phenotype matrix at 16995284.

**Supplementary file 1**

SPARQL queries to retrieve microbial data from the BacDive database, extracting NCBI genome accessions, taxonomic identifiers, and five key phenotypic traits: Gram stain status, motility, oxygen requirements, spore formation, and optimal growth temperature.

**Supplementary file 2**

SPARQL query used to query annotated genomes in the GBOL format for protein domain frequencies.

**Supplementary file 3**

Halving grid search space used for hyperparameter tuning of decision tree, random forest, gradient boosting, and XGBoost classifiers.

