## Supplementary file 1 for "MiGenPro: A linked data workflow for phenotype-genotype prediction of microbial traits using machine learning"

### Phneotype queries

#### gram.sparql

```
PREFIX fair: <http://fairbydesign.nl/ontology/>
SELECT DISTINCT (IF(CONTAINS(?accessionRaw, '.'), ?accessionRaw, CONCAT(?accessionRaw, '.1')) AS ?accession) ?gram ?taxid
WHERE {
   # Gram stain
   # Get species from genome by coupling the taxID to the NCBI taxonomy object and its taxID.
   # Get phenotype from the morphology tab.
   ?morphology fair:morphology/fair:cellMorphology/fair:`gramStain ?gram ;
   fair:sequenceInformation/fair:genomeSequences ?genome .
   ?genome fair:accession ?accessionRaw ;
         fair:database 'ncbi' ;
         fair:ncbiTaxId ?taxid ;
   FILTER( strStarts( ?accessionRaw, 'GCA_' ) ) .
}
```

#### motility.sparql

```
PREFIX fair: <http://fairbydesign.nl/ontology/>
SELECT DISTINCT (IF(CONTAINS(?accessionRaw, '.'), ?accessionRaw, CONCAT(?accessionRaw, '.1')) AS ?accession) ?motile ?taxid
where {
   # Get phenotype from the morphology tab.
   ?morphology fair:morphology/fair:cellMorphology/fair:motility ?motile ;
            fair:sequenceInformation/fair:genomeSequences ?genome .

   ?genome fair:accession ?accessionRaw ;
         fair:database 'ncbi' ;
         fair:ncbiTaxId ?taxid .
   FILTER( strStarts( ?accessionRaw, 'GCA_' ) ) .
}

```

#### oxygen.sparql

```
PREFIX fair: <http://fairbydesign.nl/ontology/>
SELECT DISTINCT (IF(CONTAINS(?accessionRaw, '.'), ?accessionRaw, CONCAT(?accessionRaw, '.1')) AS ?accession) ?oxygenResistance ?taxid
WHERE {
   # Get phenotype from the morphology tab.

   VALUES ?growth {'positive'}
```

```
    ?morphology fair:physiologyAndMetabolism/fair:oxygenTolerance/fair:oxygenTolerance ?oxygenResistance ;
                fair:sequenceInformation/fair:genomeSequences ?genome .
    ?genome fair:accession ?accessionRaw ;
            fair:database 'ncbi' ;
            fair:ncbiTaxId ?taxid ;
    FILTER(strStarts(?accessionRaw, 'GCA_')).
}
```

#### spore.sparql

```
PREFIX fair: <http://fairbydesign.nl/ontology/>
SELECT DISTINCT (IF(CONTAINS(?accessionRaw, '.'), ?accessionRaw, CONCAT(?accessionRaw, '.1')) AS ?accession) ?sporeForming ?taxid
WHERE {
    ?morphology fair:physiologyAndMetabolism/fair:sporeFormation/fair:sporeFormation ?sporeForming ;
    fair:sequenceInformation/fair:genomeSequences ?genome .
    ?genome fair:accession ?accessionRaw ;
    fair:database 'ncbi' ;
    fair:ncbiTaxId ?taxid ;
    FILTER( strStarts(?accessionRaw, 'GCA_')).
}
```

#### temperature.sparql

```
PREFIX fair: <http://fairbydesign.nl/ontology/>
SELECT DISTINCT (IF(CONTAINS(?accessionRaw, '.'), ?accessionRaw, CONCAT(?accessionRaw, '.1')) AS ?accession) ?temperatureRange ?taxid

WHERE {
    ?morphology fair:cultureAndGrowthConditions/fair:cultureTemp/fair:range  ?temperatureRange ;
                fair:cultureAndGrowthConditions/fair:cultureTemp/fair:growth ?growth ;
                fair:sequenceInformation/fair:genomeSequences ?genome .
    ?genome fair:accession ?accessionRaw ;
            fair:database 'ncbi' ;
            fair:ncbiTaxId ?taxid ;
}
```
